# The genetic architecture of local adaptation II: The QTL landscape of water-use efficiency for foxtail pine (*Pinus balfouriana* Grev. & Balf.)

**DOI:** 10.1101/038240

**Authors:** Andrew J. Eckert, Douglas E. Harwood, Brandon M. Lind, Erin M. Hobson, Annette Delfino Mix, Patricia E. Maloney, Christopher J. Friedline

## Abstract

Water availability is an important driver of the geographic distribution of many plant species, although its importance relative to other climatic variables varies across climate regimes and species. A common indirect measure of water-use efficiency (WUE) is the ratio of carbon isotopes (δ^13^C) fixed during photosynthesis, especially when analyzed in conjunction with a measure of leaf-level resource utilization (δ^15^N). Here, we test two hypotheses about the genetic architecture of WUE for foxtail pine (*Pinus balfouriana* Grev. & Balf.) using a novel mixture of double digest restriction site associated DNA sequencing, species distribution modeling, and quantitative genetics. First, we test the hypothesis that water availability is an important determinant of the geographical range of foxtail pine. Second, we test the hypothesis that variation in δ^13^C and δ^15^N is genetically based, differentiated between regional populations, and has genetic architectures that include loci of large effect. We show that precipitation-related variables structured the geographical range of foxtail pine, climate-based niches differed between regional populations, and δ^13^C and δ^15^N were heritable with moderate signals of differentiation between regional populations. A set of large-effect QTLs (*n* = 11 for δ^13^C; *n* = 10 for δ^15^N) underlying δ^13^C and δ^15^N variation, with little to no evidence of pleiotropy, was discovered using multiple-marker, half-sibling regression models. Our results represent a first approximation to the genetic architecture of these phenotypic traits, including documentation of several patterns consistent with δ^13^C being a fitness-related trait affected by natural selection.

## Introduction

Descriptions of the genetic components underlying fitness-related phenotypic variation have been a focus of quantitative genetics for over a century (Shull 1908; Fisher 1918; Mather 1941; Ford 1975; Mackay *et al*. 1994; Ritland *et al*. 2011 and references therein). These descriptions have progressed from identifications of the genetic elements affecting trait variation (e.g. Jermstad *et al*. 2001) to analysis of interactions among these elements with one another and the environment (e.g. Jermstad *et al*. 2003). Uniting all these descriptions are foundational questions about the structure, function, and evolution of genotype-phenotype maps in natural populations. For forest trees, these descriptions historically addressed traits of economic importance such as specific gravity of wood (e.g. Groover *et al*. 1994), microfibril angle (e.g. Sewell *et al*. 2000), growth (e.g. Wu 1998), and phenology (e.g. Pelgas *et al*. 2011), with the ultimate goals of marker-assisted breeding (Neale and Savolainen 2004) and trait prediction from genotypic data (Grattapaglia and Resende 2011). These traits, while economically important, often also affect fitness (especially phenology, see Sorensen 1983), so that these efforts can also be leveraged to understand the genetic basis of ecologically relevant trait variation. The linkage between traits measured in common gardens and fitness in natural populations, however, is usually assumed *post hoc,* which can lead to storytelling (Barrett and Hoekstra 2011) and oversimplification of the ecological ramifications of quantitative genetic results. Here, we address this disconnect through simultaneous use of species distribution modeling and quantitative trait locus (QTL) mapping to dissect the genetic architecture of an ecologically important phenotypic trait for foxtail pine *(Pinus balfouriana* Grev. & Balf).

The spatial and temporal distribution of all viable individuals across the Earth’s landscape for a given species is defined as its geographical range (Brown *et al*. 1996). Evolution of range sizes and structural attributes of these ranges have been studied for a variety of taxa for many decades (e.g. Mayr 1963; Antonovics 1976; Brown *et al*. 1996; Gaston 2003; Eckert *et al*. 2008; Sheth and Angert 2014). The common thread underlying these interests is the assumption that fitness of individuals within species is related to the known geographical range for each species based on the environments defined by this range, other selective pressures (i.e. competition) across this range, and the phylogeographic history that resulted in the current geographical range (Hutchinson 1957; Pulliam 2000; Chuine 2010). For example, relative fitness values within plant populations tend to be highest in their home environments and lower in novel environments at the margin or outside of known geographical ranges (reviewed in Leimu and Fischer 2008). Regardless of the relationship between this pattern and evolutionary concepts such as local adaptation, it is clear that current geographical ranges are to some degree projections of ecological niches (i.e. realized versus fundamental niches), or at least some aspect of these niches, onto geographical space (Pulliam 2000; Ettinger *et al*. 2011). Knowledge of the environmental and climatic drivers of geographical ranges can therefore be informative about links between traits responsive to these drivers and fitness.

Species distribution models (SDMs) are commonly utilized as predictive tools with which to assess the importance of environmental variables to current geographical ranges of species (Elith *et al*. 2006). At a minimum, these models are built from known occurrences of a certain species and the environmental and ecological attributes of these locations derived from either field measurements or information stored in geographical information systems (GIS) layers. Numerous approaches are available with which to build models from these data (Segurado and Araujo 2004; Elith *et al*. 2006; Phillips *et al*. 2006). Once constructed, SDMs are often used subsequently to study the evolutionary development of ranges (e.g. McCormack *et al*. 2010), as well as the effects of continued climate change on current geographical ranges (e.g. Pearson and Dawson 2003). However, there are limitations to equating SDMs, even those with good predictive abilities of current geographical ranges, with realized ecological niches and hence measures of fitness limits (Hampe 2004; Soberon and Peterson 2005; Warren and Seifert 2011). For example, individuals used to create SDMs are considered exchangeable, so that fitness variation among individuals is ignored (Hampe 2004). Some of these issues, especially those related to exchangeability of individuals within species, can be addressed through a careful matching of modeling units (e.g. genetically differentiated populations within species; *sensu* Davis *et al*. 2005), geographical scale (e.g. the geographical scale relevant to the genetically differentiated populations), and the research questions of interest.

Water is crucial to the survival of many plant species (e.g. Sorenson 1983), although its importance relative to other environmental factors varies depending upon the environmental factors that are most limiting within local environments (Dudley 1996). The intrinsic efficiency by which plants use water (WUE) is defined as the ratio of net assimilation of carbon from CO_2_ during photosynthesis to the loss of water during transpiration (Bacon 2004). Carbon isotopic composition (δ^13^C) is an indirect measure of intrinsic WUE and is based upon the ratio of two isotopes of carbon (^13^C and ^12^C) within plant tissue standardized to a reference. This ratio is related to WUE because it has been demonstrated that the discrimination by C_3_ plants of ^13^CO_2_ relative to ^12^CO_2_ is correlated to the ratio of carbon assimilation during photosynthesis to stomatal conductance (Farquhar *et al*. 1982; Farquhar and Richards 1984; e.g. Zhang and Marshall 1994). The physiological and environmental mechanisms, however, driving the linkage between δ^13^C and intrinsic WUE at various levels of biological organization are numerous, so that the expected linear relationship between δ^13^C and WUE may not always hold (Seibt *et al*. 2008). For example, differences in δ^13^C across individual plants at the leaf level can result from changes in carbon to nitrogen allocation during carboxylation, variation in leaf structure and morphology, and/or variation in available CO_2_ (Seibt *et al*. 2008). Within a common environment, however, it is assumed that variation in available amounts of atmospheric CO_2_ is negligible. Variation for δ^13^C across individual plants in these common environments should therefore reflect variation for intrinsic WUE. Indeed, previous research in conifers has established that variation in δ^13^C across individual plants is heritable (Seiler and Johnson 1988; Cregg 1993; Brendel *et al*. 2002; Baltunis *et al*. 2008; Cumbie *et al*. 2011), is polygenic, yet comprised of a mixture of large and small effect loci (Brendel *et al*. 2002; Gonzalez-Martinez *et al*. 2008; Cumbie *et al*. 2011; Marguerit *et al*. 2014), and that it often reflects variation for intrinsic WUE through leaf level assimilation (Zhang and Marshall 1994; Brendel *et al*. 2002; Cumbie *et al*. 2011; Marguerit *et al*. 2014).

Water availability is often an important driver of tree distributions (Stephenson 1990 and references therein), especially in Mediterranean climates (e.g. Baldocchi and Xu 2007; Lutz *et al*. 2010). This importance is evident through increased tree mortality as a function of both direct and indirect consequences associated with changing water availability (van Mantgem *et al*. 2009; Allen *et al*. 2010). Regional and local water availability will likely be altered, either through changes to annual precipitation totals or the seasonality of precipitation, under most climate change scenarios, especially in ecosystems dependent on residual summer snow-packs (Barnett *et al*. 2005). The ability of natural populations of forest trees to respond to changing water availability is linked to segregating genetic variation for traits responsive to water availability (Aitken *et al*. 2008). Knowledge of the genetic architecture of such traits, therefore, provides an important resource for assessing forest health, as well as the genetics of adaptation (Neale and Kremer 2011). Here, we test two hypotheses about the genetic architecture of WUE for foxtail pine – (i) water availability is an important determinant of the geographical range of foxtail pine and hence fitness and (ii) variation in δ^13^C and δ^15^N is genetically based, differentiated between regional populations, and has genetic architectures that include loci of large effect. We subsequently discuss how the integration of results from disparate fields of research (i.e. genomics, ecology, and quantitative genetics) provides information useful to foundational tests about the genetic architecture of local adaptation and its evolution (*cf*. Friedline *et al*. 2015).

## Materials and Methods

### Focal Species

Foxtail pine is one of three species classified within subsection *Balfourianae* of section *Parrya* within subgenus *Strobus*. It is generally regarded as the sister taxon to Great Basin bristlecone pine (*P. longaeva* D. K. Bailey; see Eckert and Hall 2006). The distribution of this species is relegated to the high elevation mountains of California, with all known occurrences being in either the Klamath Mountains of northern California or in the high elevations of the southern Sierra Nevada (Figure S1). These two regions are separated by approximately 500 km and differ in climate, soils, and forest composition (Ornduff 1974; Eckert and Sawyer 2002; Barbour *et al*. 2007).

### Common Garden

A common garden representing 141 maternal foxtail pine trees was established at the Institute of Forest Genetics (Placerville, CA) during 2011 and 2012 using a randomized block design. Cones were collected from 141 maternal trees sampled range-wide, with 72 sampled from the Klamath Mountains and 69 from the southern Sierra Nevada region. For each maternal tree, 35 – 100 seeds were germinated and grown in standard conditions as outlined in Eckert *et al*. (2015). More information about the common garden can be obtained from Friedline *et al*. (2015). Of these 141 maternal trees, offspring, assumed to be half-siblings, from five were selected for analysis of water-use efficiency (see **Phenotype determination**, Table 1). The megagametophyte associated with each germinated seed from these five maternal trees was rescued and used to construct a high-density linkage map based on four of the five maternal trees (Friedline *et al*. 2015). The seedlings from each maternal tree were allowed to grow for a full year after which needles were sampled (*n* = 32 to 40/maternal tree) for determination of phenotypes and genotypes. As done by Friedline *et al*. (2015), families were named using colors (i.e. these were the colors of family identifier tags in the common garden), with families sampled from the Klamath Mountains being labeled as blue, yellow, and purple and families sampled from the southern Sierra Nevada being labeled as red and green.

**Table 1.** Summary of the families (*n* = 5) used for QTL mapping

|  | Red | Green | Purple | Blue | Yellow |
| --- | --- | --- | --- | --- | --- |
| Latitude | 36.448075 | 36.448075 | 41.319871 | 41.195910 | 41.748267 |
| Longitude | -118.170644 | -118.170644 | -122.479184 | -122.792240 | -123.133233 |
| Elevation (m) | 3352.80 | 3352.80 | 2397.56 | 2103.12 | 2103.12 |
| Siblings<sup>a</sup> | 35 | 40 | 34 | 40 | 32 |
| Locality | Cottonwood | Cottonwood | Mt. Eddy | East Boulder | Lake |
|  | Pass | Pass |  | Lake | Mountain |
| Region | SN | SN | KM | KM | KM |

<sup>a</sup>These counts represent the numbers of siblings genotyped and phenotyped for each family. Additional siblings for each family are still growing within the common garden (see **Materials and Methods**).

### Phenotype Determination

Two phenotypic traits were measured from needle tissue sampled from each growing seedling – carbon isotope discrimination (δ^13^C) and foliar nitrogen content (δ^15^N). These were chosen because (δ^13^C) is a proxy for intrinsic WUE (Farquhar *et al*. 1982; Farquhar and Richards 1984), while δ^15^N is a proxy for plant growth and resource utilization during photosynthesis (Prasolova *et al*. 2000). Tissue was sampled in year 1 of growth, which was also prior to formation of randomized blocks in the common garden. Given the age of the seedlings, sampling of enough needle tissue for determination of phenotypes and genotypes was destructive. Thus, only a subset of the seedlings per maternal tree was used. For these seedlings, all available needles were sampled, cleaned and separated into those used for genotype determination and those used for phenotype determination. For phenotype determination, needles were placed into a mortar with liquid nitrogen and coarsely ground by hand using a pestle. The resulting needle tissue was then transferred into 20 ml glass vials and oven-dried at 60°C for 96 hrs. Approximately, 2 to 3 mg of ground and dried needle tissue from each seedling was subsequently placed into individual wells comprising a 96 well microtiter plate. Samples were analyzed for δ^13^C and δ^15^N at the Stable Isotope Facility at UC Davis (http://stableisotopefacility.ucdavis.edu/). Data are presented as carbon isotope ratios for δ^13^C (‰) and weight for δ^15^N (µg).

### Sequence Analysis and Genotype Determination

Total genomic DNA was extracted from the remaining needles from each sampled seedling using Qiagen DNeasy 96 Plant Kits following the manufacturer’s protocol. The resulting total genomic DNA for each seedling was quantified using spectrophotometry as implemented with a Thermo Scientific NanoDrop 8000. Following quantification, samples were prepared for double digest restriction site associated DNA sequencing (ddRADseq) following the protocols of Parchman *et al*. (2012) as implemented for foxtail pine by Friedline *et al*. (2015). All samples had concentrations of total genomic DNA in the range of 15 to 60 ng/ul. In brief, this protocol proceeds via restriction digests of total genomic DNA for each sample using EcoR1 and Mse1, ligation of adapters that include the Illumina primer, universal M13 primers, and 8 - 10 bp barcodes, PCR amplification, and size selection of the PCR amplified and ligated restriction digests. In our protocol, multiplexing (i.e. pooling) occurred post PCR and size selection was carried out using 1.0% agarose gels run for 1 hour at 110 volts in 1X TAE buffer. All data are based on sequencing fragments in the size range of 300 to 500 bp on the Illumina HiSeq 2500. DNA sequencing was performed at the VCU Nucleic Acid Research Facility (http://www.narf.vcu.edu/index.html).

Raw FASTQ sequences were quality-checked and filtered as in Friedline *et al*. (2015). Briefly, reads must pass a three-stage filtering procedure to be retained for downstream analysis. First, if the average quality for all bases in the read was below 30, the read was discarded. Second, a five-base pair sliding window was evaluated along each raw sequence. Consecutive windows were retained if their mean quality was greater-than or equal-to 30. If the mean score of a window fell below this threshold, the read was trimmed at this point. If the length after trimming was at least 50% of the original read length, the read was kept, otherwise it was discarded. Finally, if 20% of the bases in the original read had quality scores below 30, the entire read was discarded, even if its average quality met the inclusion threshold. The reads that passed quality filtering were demultiplexed and assigned to individual trees in one of five families: Blue, Green, Purple, Red, or Yellow.

Sequences were aligned to the linkage map assembly (Friedline *et al*. 2015) and read groups were added using Bowtie2 version 2.2.4 (Langmead and Salzberg 2012) using the - very-sensitive-local set of options. Each alignment was checked and marked for PCR artifacts using Picard (http://picard.sourceforge.net, svn 03a1d72). Variants were called using the mulitallelic caller from samtools version 1.1 (Li *et al*. 2009), specifying diploidy for all individuals. The resulting VCF file was processed using VCFtools version 0.1.12.b (Danecek *et al*. 2011), retaining only bialleleic SNPs that mapped to positions on the linkage map defined in Friedline *et. al* (2015) with quality (-- minQ) of at least 20. All read processing and variant calling pipeline code, Python 3.4.3 and R version 3.2.0 (R Core Team 2015), can be found as IPython (Pérez and Granger 2007) notebooks and associated files at http://www.github.com/cfriedline/foxtailwue.

Once genotypes were called for all loci on the linkage map of Friedline *et al*. (2015), we selected one SNP per position on the linkage map based on minimizing the amount of missing data and being polymorphic in the most families. Missing genotype data were subsequently imputed for each linkage group using the default settings of the program fastPHASE ver. 1.2 (Scheet and Stephens 2006), with families used as populations. To account for uncertainty in genotype imputation, we estimated posterior probabilities of each possible genotype (i.e. 0, 1, or 2) at each locus using 1,000 haplotype reconstructions provided by fastPHASE, which were used subsequently used as weights in a weighted average of the minor allele count. These weighted averages were then rounded to the closest value (0, 1, or 2) following normal rounding rules (i.e. round downward if the tenths position is less than five, otherwise round up).

### Species Distribution Modeling

We used species distribution models (SDMs) to justify water-use efficiency as a fitness-related trait and to quantify niches of each regional population relative to one another. The former provides an *a priori* justification for the measured traits as ecologically relevant, while the latter provides an estimate of niche differentiation between regional populations comparable to the effect of region on trait differentiation (see **Quantitative Genetic Analysis**).

Species distribution models were used to assess the relative importance of precipitation-related and temperature-related variables to the distribution of foxtail pine. We utilized the approach of maximum entropy (MaxEnt; Phillips *et al*. 2006) to construct SDMs. Known locations of foxtail pine within each regional population *(n =* 93 Klamath Mountains, *n =* 207 southern Sierra Nevada) were gathered from digitized herbarium records available through the Jepson Herbarium located at the University of California, Berkeley (http://ucjeps.berkeley.edu/). When the latitude and longitude of locations associated with these herbarium records were missing, visual inspections of maps from Google Earth were used to find the best approximation to the locality described on the herbarium sample. Climate data for each regional population were obtained from WordClim (http://www.worldclim.org/) and are represented as 19 bioclimatic variables, which are functions of temperature and precipitation variables (Table S1), given at a resolution of 30 arc-seconds (∼1 km). The generic grid files available from the WorldClim website were trimmed for each climate variable using the *raster* library in R and the following geographical extent: minimum longitude: −124.0°, maximum longitude: −117.5°, minimum latitude: 35.0°, maximum latitude: 42.5°. Using these trimmed grid files and the location information pruned of duplicate observations (*n*_pruned_ = 65 Klamath Mountains, *n*_pruned_ = 144 southern Sierra Nevada), the MaxEnt software version 3.3.3k (https://www.cs.princeton.edu/∼schapire/maxent/) was used to build a SDM for each regional population. MaxEnt was run using the cross-validation option for model assessment, 10 replicates, a maximum number of background points of 10,000, and jackknife analysis to evaluate variable importance. Measures of variable importance (i.e. variable contribution and permutation importance scores) and the results of the jackknife analyses were used to assess the relative roles of temperature-related and precipitation-related variables to each SDM.

We used also used SDMs to quantify niche differentiation between regional populations of foxtail pine (Warren *et al*. 2008). We tested two null hypotheses. First, we tested the null hypothesis that the two SDMs were based on a single, underlying SDM common to each regional population. Second, we tested the null hypothesis that the two SDMs are no more differentiated than those randomly drawn from a common SDM with non-overlapping geographical distributions for each regional population. Both tests are based on the *D* and *I* statistics given by Warren *et al*. (2008). The former null hypothesis was tested using the *niche.equivalency.test* function in the *phyloclim* library in R, while the latter null hypothesis was tested using the *bg.similarity.test* function in the same R library. Both tests were based on *n =* 100 permutations to derive null distributions of test statistics.

### Quantitative Genetic Analysis

We performed two sets of analyses to dissect the genetic basis of water-use efficiency for foxtail pine. First, we demonstrated that variation for the measured traits was genetically based using standard methods to decompose trait variance into effects of families, regions, and environment (Lynch and Walsh 1998). Second, we fit single and multiple QTL models to dissect the genetic basis of each trait into their genetic components using the regression methods of Knott *et al*. (1996).

The genetic basis for each measured trait was assessed using linear models. We fit three different linear models to the observed data for each trait: (1) a fixed effect model containing only a grand mean (i.e. intercept), (2) a linear mixed model with a grand mean as a fixed effect plus a random effect of family, and (3) a linear mixed model of a grand mean as a fixed effect plus a random effect of region plus a random effect of family nested within region. Uncertainty in parameter estimates from each model was assessed using parametric bootstrapping *(n =* 1,000 replicated simulations) as carried out with the *simulate* function in R. Models were compared using the Akaike Information Criterion (AIC), with Akaike weights used to assess the conditional probabilities for each model (Burnham and Anderson 2002). If models containing random effects for families or models containing random effects for regions and families nested within regions fit the data better than a model with only a grand mean, then we concluded that there were non-zero heritabilities for these traits. If we assume that all offspring within each family were half-siblings, we could estimate narrow-sense heritability as *h^2^ =* 4σ^2^_fam_/(σ^2^f_am_ + σ^2^res), where σ^2^f_am_ is the variance due to family nested within region and σ^2^_res_ is the residual variance. Given the small number of families, however, we avoided this estimation, as we were interested only in detecting non-zero heritability and not precise estimation of its magnitude. Linear models with fixed effects were fit using the *lm* function, while linear mixed models were fit using maximum likelihood as employed in the *lmer* function of the *lme4* library of R. Log-likelihood and AIC values were extracted for each fitted model using the *logLik* and *AIC* functions in R, respectively.

The genetic basis of each trait was dissected using the least squares regression approach of Knott *et al*. (1996) for outbred, half-sibling families, where probabilities of allelic inheritance due to the common parent were used as predictors for each trait. Significance of the regression model was determined using a *F*-test calculated at 1-cM intervals, with the distribution of this statistic under a null model of no QTLs generated via a permutation scheme (Churchill and Doerge 1994). The common parent in our analyses was the maternal tree, we assumed that all offspring per maternal tree were half-siblings, and we used 1,000 permutations to generate null distributions of *F*-statistics. Permutations were used to create null distributions for *F*-statistics at the level of the entire genome (i.e. all linkage groups) and for each chromosome (i.e. linkage group) separately. We initially fit models of one QTL per linkage group using three significance thresholds: (1) α = 0.05 at the level of the entire genome (major QTL), (2) α = 0.01 at the level of a particular chromosome (minor QTL), and (3) α = 0.05 at the level of a particular chromosome (suggestive QTL). For each QTL, we estimated the percent variance explained (PVE) as PVE = 4[1 − (MSE_full_/MSE_reduced_)], where MSE_full_ and M S Educed are the mean square errors of the full and reduced models, respectively *(cf*. Everett and Seeb 2014). Following Knott *et al*. (1996), estimates of PVE were scaled by (1 − *2r*)^2^, where *r* is the recombination frequency between the marker and QTL (i.e. *r =* 0.01 for a 1-cM scan of each linkage group). Uncertainty in the position of the QTL was assessed using bootstrapping *(n =* 1,000 replicates). For each linkage group with a statistically significant QTL, we subsequently fit a model of two QTLs using the same approach, with the only differences being the use of asymptotic null distributions to test the statistical significance of the observed *F*-statistics and the lack of adjustments to estimates of the PVE for multiple QTL models. All analyses were conducted with the HSportlets module on GridQTL ver. 3.3.0 (Seaton *et al*. 2006; Allen *et al*. 2012) using the linkage map for foxtail pine reported by Friedline *et al*. (2015).

## Results

### Sequence Analysis and Genotype Determination

From two lanes of HiSeq sequencing, we obtained 148,685,598 and 160,770,417 reads from lane 1 (length = 101 bp, %GC = 40) and lane 2 (length = 101 bp, %GC = 41), respectively. Following read filtering, we retained 77,568,370 (length = 49 − 101 bp, %GC = 40) reads from lane 1 and 107,372,313 (length = 49 − 101, %GC = 40) reads from lane 2. A summary of the sequencing output and quality can be found in Table 2. The highest quality and most reads came from the Blue and Red families, while the Green family produced the smallest number of reads. Similarly, the Blue and Red families had the highest percentages of reads mapping to the assembly. The quality of reads across all families was sufficiently high, with average quality of any base of approximately 38. Graphical summaries of missing data and quality metrics are available in Figures S2 and S3. We filtered SNPs at the same position on the linkage map down to a set of 843 loci with the least amount of missing data and polymorphism in the most families. At these 843 SNPs, missing data averaged 58.0% (0% − 95.6%). Missing data were subsequently imputed using the marker ordering from Friedline *et al*. (2015) and fastPHASE.

**Table 2.**
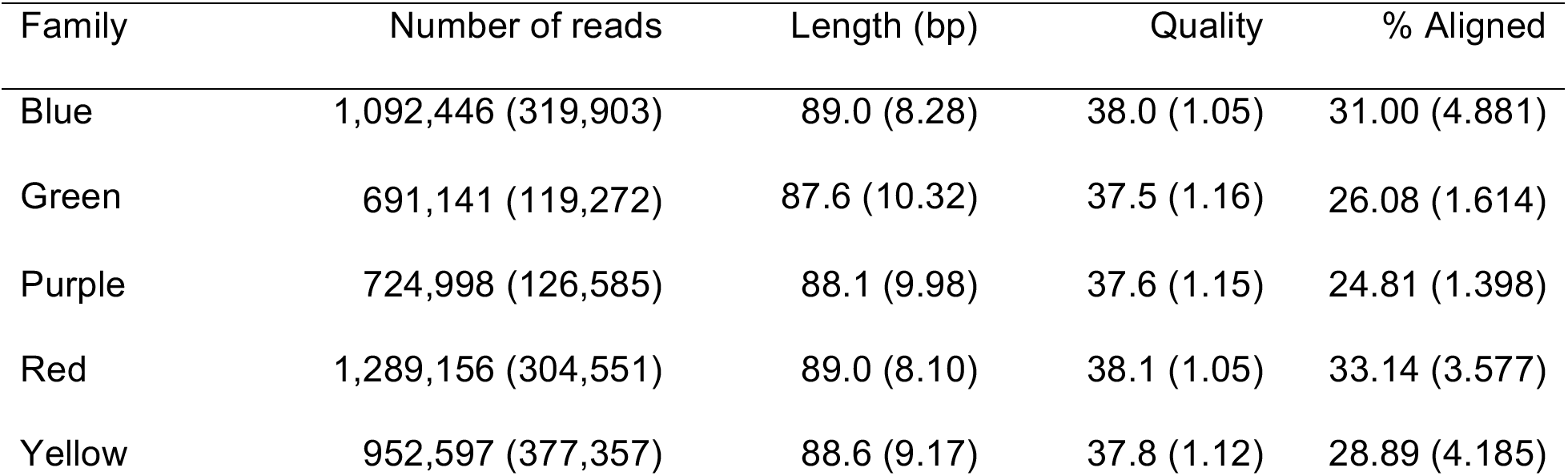
Mean and standard deviation (in parentheses) of read metrics by family

### Species Distribution Modeling

Species distribution models were good predictors of the current geographical ranges for each regional population of foxtail pine (Figures 1, S1). Estimates of the area under the receiver operating characteristic curves (ROC curves) were near 1.0 for each model for both the training and test set of samples (Figure S4). Exceptions to this pattern included low to moderate probabilities of occurrence outside the current geographical distribution for the Klamath Mountains, which were centered on the northern Sierra Nevada, and a slightly expanded range north and south of the known range limits in the southern Sierra Nevada. Foxtail pine is known to be absent from these regions. In both cases, the probabilities of occurrence were less, often much less, than 0.40. The SDM based on the Klamath Mountains predicted a near zero probability for cells within the range of the southern Sierra Nevada and vice versa.

Foxtail pine inhabits the cooler portions of each region in which it is currently located (Figures S5 - S6). For precipitation-related variables, however, foxtail pine in the Klamath Mountains inhabits slightly wetter localities relative to background localities, while in the southern Sierra Nevada foxtail pine inhabits drier localities relative to background localities. The climates inhabited by foxtail pine in each region also differ. In general, differences between the climates inhabited by each regional population were consistent with the Klamath Mountains being warmer, yet less variable in temperature throughout the year, and wetter, yet slightly more variable in precipitation throughout the year, relative to the southern Sierra Nevada. For example, mean annual precipitation was almost twice as high in the Klamath Mountains as in the southern Sierra Nevada (1179.66 mm versus 650.03 mm, respectively), yet the distribution of precipitation was slightly more variable throughout the year (e.g. precipitation of the driest month: 11.78 mm versus 12.41 mm, respectively; coefficient of variation across months: 65.86 versus 65.02, respectively).

Bioclimatic variables used to predict occurrences of foxtail pine within each regional population were highly correlated with one another (Figure S7). Sets of correlated variables are difficult to evaluate as contributing to SDMs (Warren and Seifert 2011). We, therefore, used several different measures of variable importance. Inspection of variable contribution scores revealed that temperature-related and precipitation-related variables were differentially important across SDMs for each region (Figure 1; Table S2). Temperature-related variables, specifically mean diurnal range (Bio2), isothermality (Bio3), and maximum temperature of the warmest month (Bio5), were most important for the southern Sierra Nevada population, whereas precipitation-related variables, specifically precipitation of the driest quarter (Bio17) and precipitation of the wettest quarter (Bio16), were most important for the Klamath Mountains population. This pattern, however, was reversed when using permutation importance scores, despite a moderate correlation between rankings of importance based on variable contribution and permutation importance scores (Figures 2, S8; Table S4). Temperature-related variables became more important for the Klamath Mountains, specifically annual temperature (Bio1), while precipitation-related variables became more important for the southern Sierra Nevada population, specifically precipitation seasonality (Bio15) and mean temperature of the wettest quarter (Bio8). Jackknife analysis of variable importance based on AUC, test gain, and regularized test gain, however, were consistent with both temperature-related and precipitation-related variables as being important for the Klamath Mountains population (Figures S9 - S11). For example, mean annual temperature (Bio1), maximum temperature of the warmest quarter (Bio5), mean temperature of the driest quarter (Bio9), mean temperature of the warmest quarter (Bio10), precipitation of the driest quarter (Bio17), and precipitation of the warmest quarter (Bio18) all contributed significantly to the SDM for the Klamath Mountains population (Figure S11), although no one variable contained much information that was not present in at least one of the others. In contrast, jackknife analysis of variable importance based on AUC, test gain, and regularized test gain were consistent with primarily temperature-related variables, specifically mean annual temperature (Bio1), mean diurnal range (Bio2), maximum temperature of the warmest month (Bio5), and the mean temperature of the warmest quarter (Bio10), driving the SDM for the southern Sierra Nevada population (Figures S12 - S14). As with the SDM for the Klamath Mountains population, however, no one variable contained information that was not present in at least one of the others (Figure S14).

**Figure 1.**
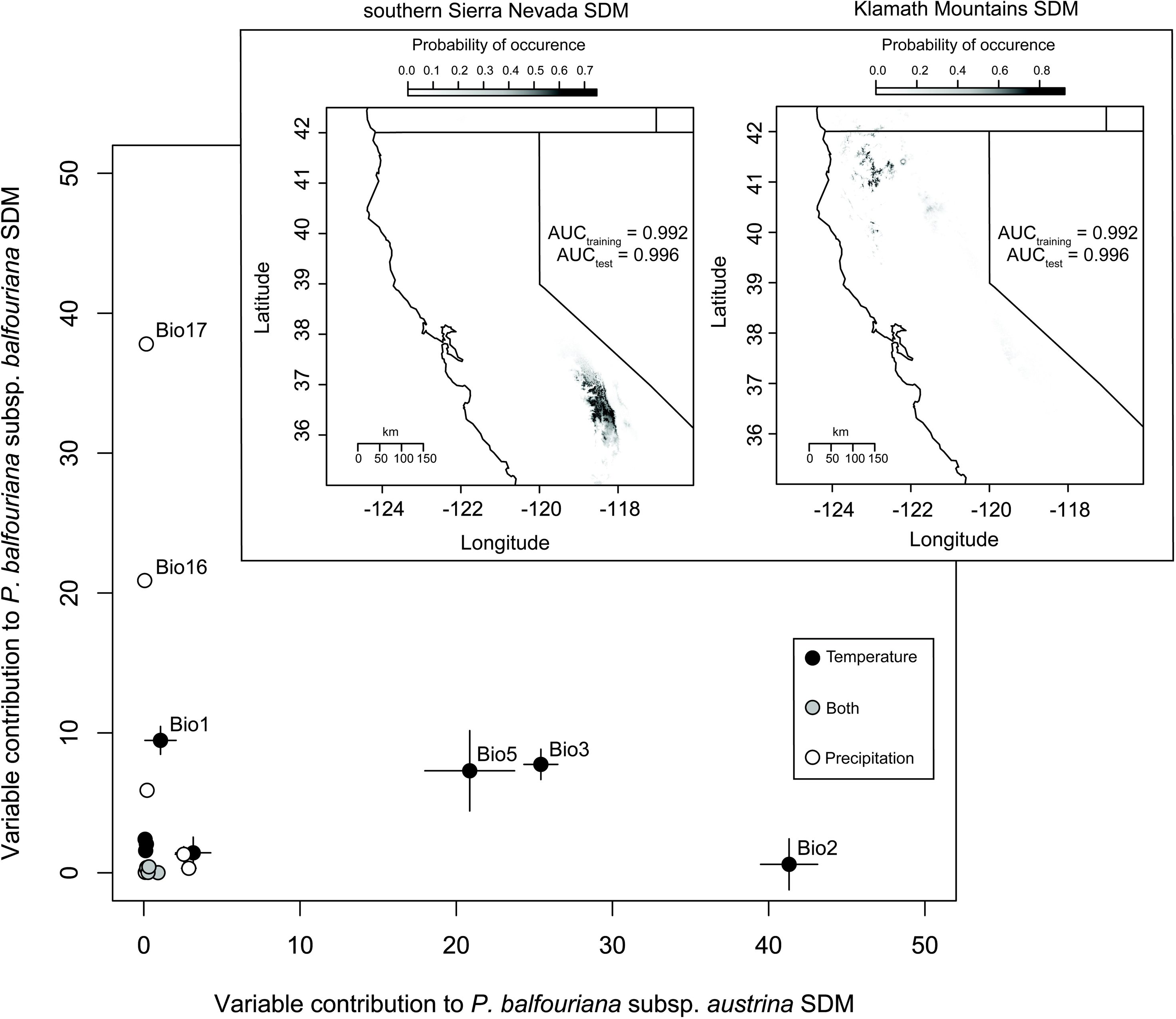
Species distribution models (SDMs) created using MaxEnt are good predictors of the current geographical range of foxtail pine (inlaid maps; AUC = area under the receiver operating characteristic curve). Precipitation and temperature-related variables are differentially important, as measured by variable contributions to each model, to the SDM of each regional population of foxtail pine, with precipitation-related variables more important for the Klamath Region and temperature-related variables more important for the southern Sierra Nevada. Variable contribution scores (+/-1 standard deviation derived from 10 replicated runs of MaxEnt per SDM) are uncorrelated (Spearman’s ρ = −0.065). For symbols without apparent error bars, the diameter of the circle was greater than the standard deviation.

Predicted niches based on SDMs for each regional population were dissimilar, with estimates of *D* (0.072) and *I* (0.258) being much closer to zero (dissimilar) than to 1 (similar) (Figure S15). These differences were significant enough to reject a null model of a single shared SDM common to both regional populations *(P <* 0.01 for *D* and *I*). Even if differences were accounted for in the background environments of each regional population (Figure S5), the predicted niches were statistically different (*P* < 0.05 for both *D* and *I*). Replicating the analyses for climate variables related only to temperature or only to precipitation revealed that niche divergence was stronger for precipitation-related variables (*D*_precip_ = 0.074; *I*_precip_ = 0.271) relative to temperature-related variables (*D*_temp_ = 0.124, *I*_temp_ = 0.376). Therefore, regional populations of foxtail pine have divergent climatic niches, with precipitation-related variables more differentiated than temperature-related variables.

### Quantitative Genetic Analysis

Variation across siblings measured within the common garden was genetically based for each trait (Table 3). Family identifiers nested within regional populations accounted for sizeable portions of the total variance for δ^13^C (σ^2^_fam_/[σ^2^_reg_+ σ^2^_fam_+ σ^2^_res_] = 24.76%) and δ^15^N (σ^2^_fam_/[σ^2^_reg_+ σ^2^_fam_+ σ^2^_res_] = 24.45%). This was consistent with the differences among predicted family means for both traits (Figure 2), which were positively correlated (Figure 3), but not significantly so (Pearson’s *r* = 0.415; *P* = 0.487). Regional identifiers, however, were differentially important across traits, with these identifiers accounting for marginally more variance than family identifiers for δ^13^C (26.01%) but less than 10% of the total variance for δ^15^N (Figure 2). The joint effect of family and regional identifiers (i.e. the total genetic effect = [σ^2^_reg_ + σ^2^_fam_]/[σ^2^_reg_+ σ^2^_fam_+ σ^2^_res_]), however, was large for each trait (δ^13^C: 50.78%; δ^15^N: 29.75%). Comparisons of linear models progressing from intercept only to an intercept plus families nested within regions using AIC, revealed that a linear mixed model with an intercept and families was the best fit (AIC = 310.29 for δ^13^C; AIC = 1031.26 for δ^15^N; Table 4). Comparison to other models using AIC weights, however, revealed that the most complex model of an intercept plus region plus families nested within regions had a reasonably high conditional probability (AIC weight = 0.36 δ^13^C; AIC weight = 0.28 for δ^15^N; Table 4) relative to those for the best model (δ^13^C = 0.64; δ^15^N = 0.72) for each phenotypic trait.

**Figure 2.**
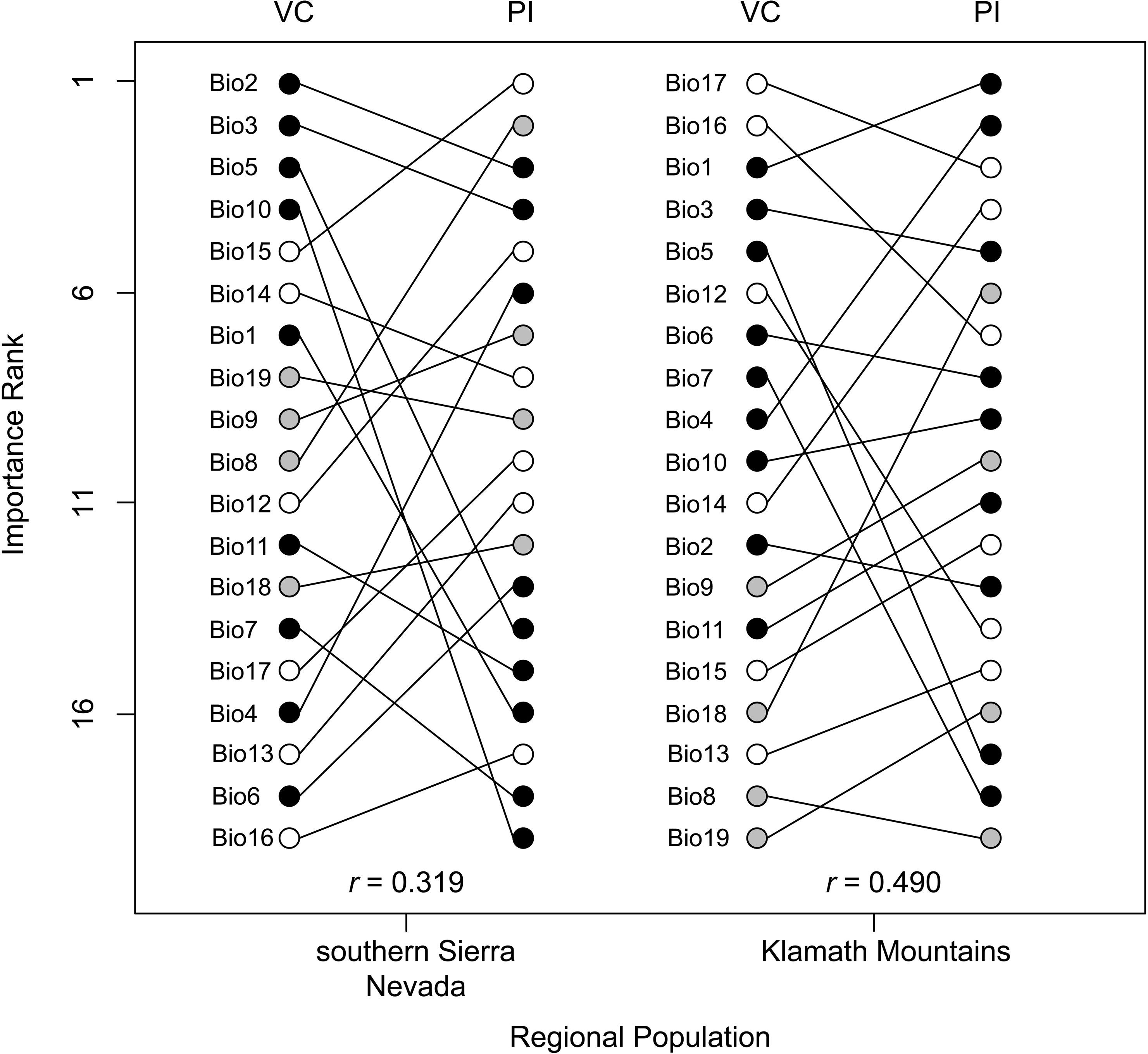
Ranks of variable importance (low rank = more important) based on variable contribution (VC) scores and permutation importance (PI) scores to the SDM for each regional population are moderately correlated *(r =* Spearman’s ρ). Variable types are denoted using filled circles, with black used for temperature-related variables, white for precipitation-related variables, and gray for variables related to both temperature and precipitation.

**Figure 3.**
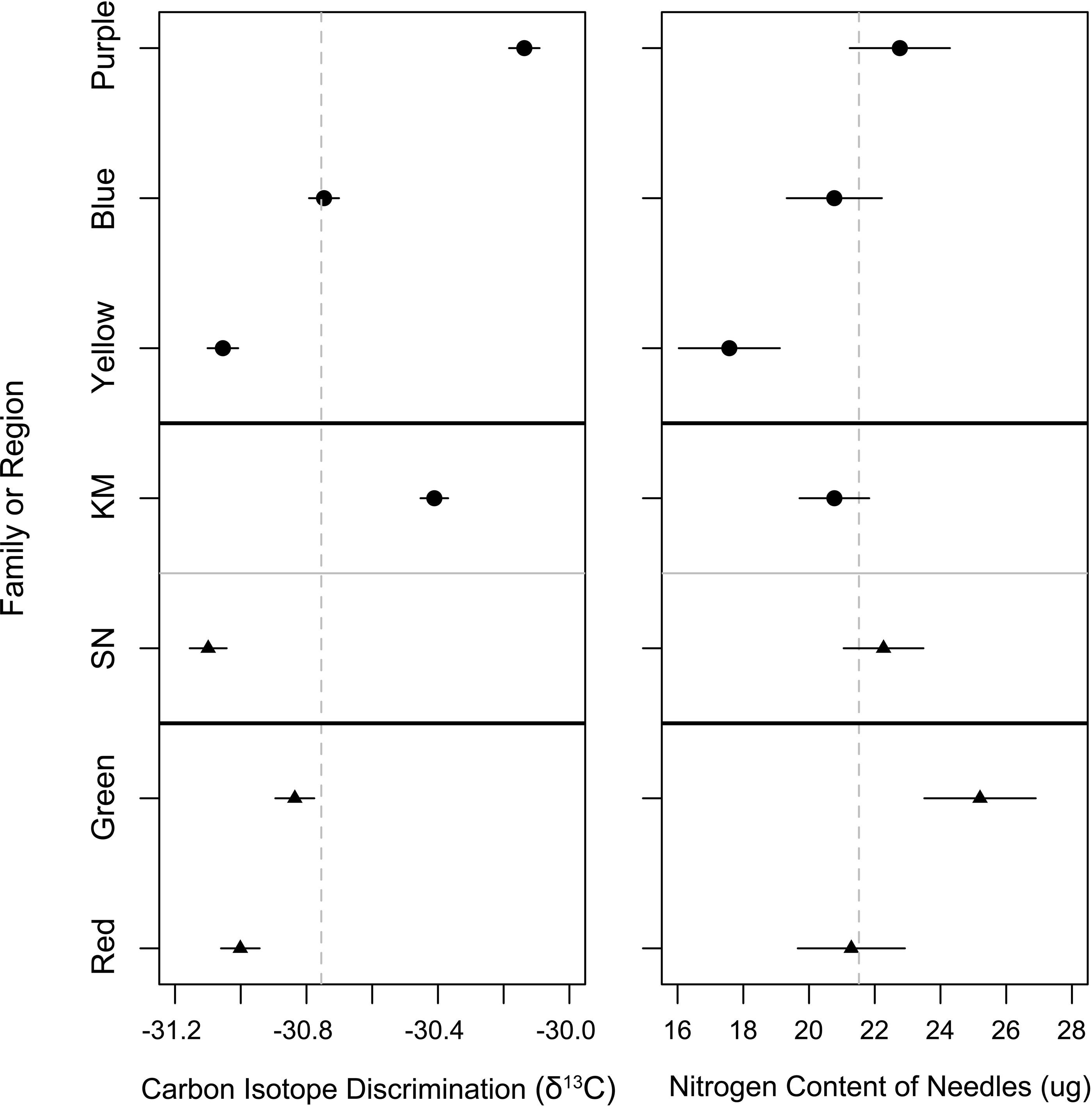
Familial and regional level means (+/-1 standard error) by trait (left: δ^13^C, right: δ^15^N) are differentiated across families and regions relative to the global mean. Dashed gray lines give global means across all families for each trait. Estimates for the Klamath Mountains (KM) are given as filled circles, while estimates for the southern Sierra Nevada (SN) are given as filled triangles. Familial names are given as colors (see **Materials and Methods**).

**Table 3.** Attributes of linear mixed models used to estimate familial and regional effects for each phenotypic trait. Values in parentheses are 95% parametric bootstrap confidence intervals (see **Materials and Methods**).

| Model Attribute | $\delta^{13}\text{C}$ | $\delta^{15}\text{N}$ |
| --- | --- | --- |
| $\log L$ | -151.705<br>(-167.029 – -130.520) | -512.587<br>(-528.588 – -491.444) |
| Intercept | -30.755<br>(-31.439 – -30.075) | 21.519<br>(18.615 – 24.596) |
| Family variance component ($\sigma^2_{\text{fam}}$) | 0.159<br>(0.002 – 0.432) | 7.826<br>(0.000 – 17.521) |
| Region variance component ($\sigma^2_{\text{reg}}$) | 0.167<br>(0.000 – 0.538) | 1.696<br>(0.000 – 9.933) |
| Residual variance component ($\sigma^2_{\text{res}}$) | 0.316<br>(0.249 – 0.384) | 22.486<br>(17.927 – 27.912) |

**Table 4.** Comparisons of linear mixed models using the Akaike Information Criterion (AIC) by trait were used to select the best model (bolded text). In these models, the intercept was a fixed effect, while families nested within regions and regions were random effects.

| Model | $\delta^{13}\text{C}$ | | $\delta^{15}\text{N}$ | |
| --- | --- | --- | --- | --- |
|  | AIC | AIC weight<sup>a</sup> | AIC | AIC weight<sup>a</sup> |
| Intercept | 408.10 | $3.66 \times 10^{-22}$ | 1071.76 | $1.16 \times 10^{-9}$ |
| **Intercept + family** | **310.29** | **0.64** | **1031.26** | **0.72** |
| Intercept + family + region | 311.41 | 0.36 | 1033.17 | 0.28 |

<sup>a</sup>The AIC weight is calculated using the standardized relative likelihoods, where the relative likelihood is given as $e^{(-0.5 \times \Delta\text{AIC})}$ . For this calculation, $\Delta\text{AIC}$ is the difference between the AIC for each model and the AIC for the best model (bolded text), where the best model is the one with the lowest AIC. The weights are then calculated as each of relative likelihoods over the sum of the relative likelihoods, thus making the sum of the weights equal to 1. Akaike weights can also be considered as the conditional probabilities for each model.

We dissected the genetic basis of the heritable variation evident for each trait from the linear mixed model analysis using the regression-based approach to QTL mapping of Knott *et al*. (1996). Application of one-locus models (i.e. a maximum of one-locus per linkage group) resulted in a set of 11 QTLs across all linkage groups and both traits (Table 5; Figure 4). For δ^13^C, six QTLs were discovered, with two discovered at the most stringent significance level (genome-wide permutation-based α = 0.05) and four at the least stringent significance level (linkage group specific permutation-based α = 0.05). Effect sizes for these QTLs were large to moderate, with the percent variation explained (PVE) ranging from 47.807% to 24.066%. For δ^15^N, five QTLs were discovered, with one QTL at the most stringent significance level, two at the intermediate significance level (linkage group specific permutation-based α = 0.01), and two at the least stringent significance level. Effect sizes for these QTLs were also large to moderate, with PVE varying from 39.773% to 25.058%. There was moderate autocorrelation for the *F*-statistic at a resolution of 6 cM or less for δ^13^C and 3 cM or less δ^15^N (Figure S16), but there was no correlation between *F*-statistics for each trait (Pearson’s *r*: −0.014, *P* = 0.734; Figure S17). In general, 95% confidence levels of positions for each QTL were large (Table 5).

**Figure 4.**
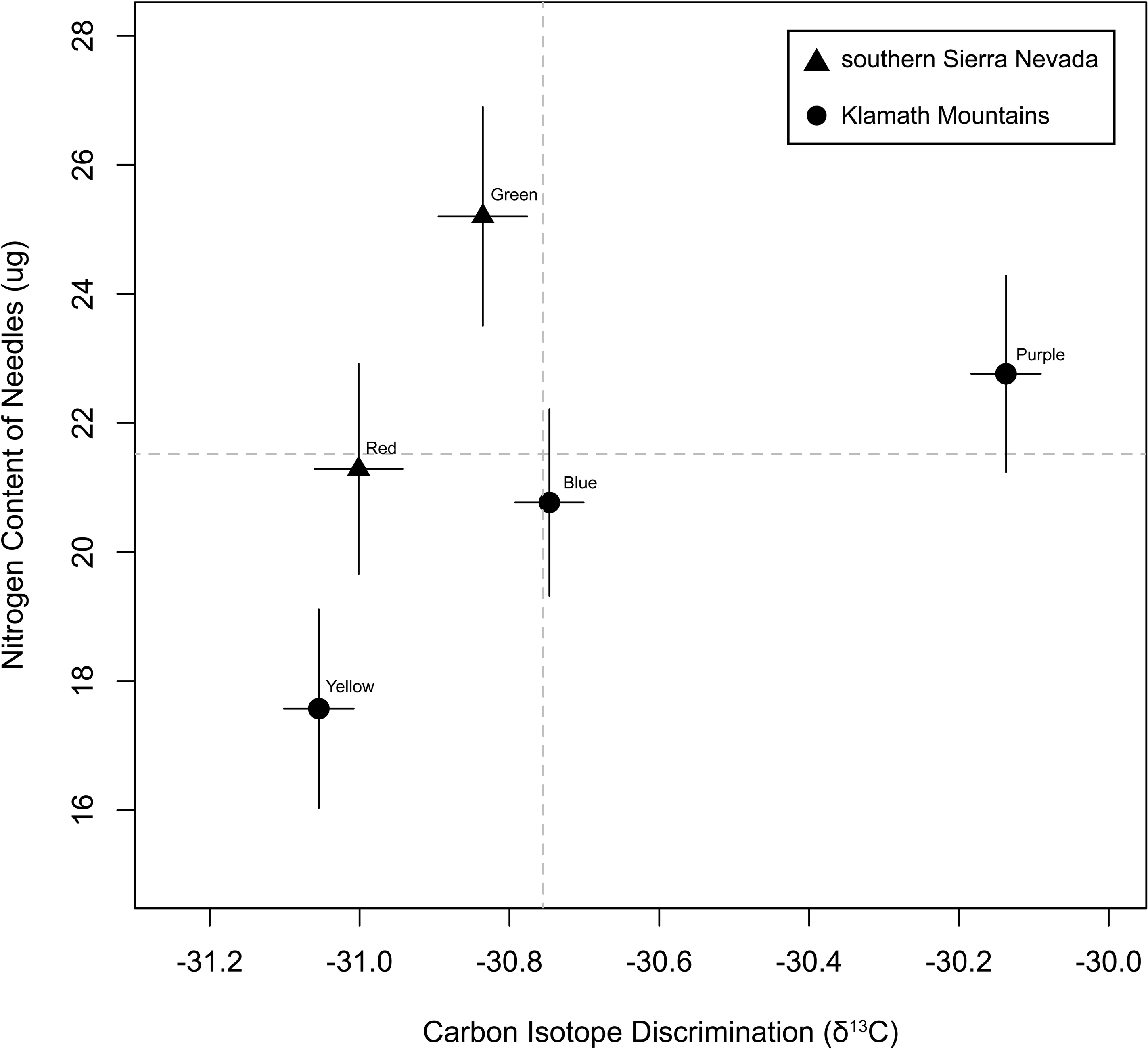
The relationship between traits based on family means (+/-1 standard error) is positive (Pearson’s *r =* 0.415), although statistically non-significant at α = 0.05 *(P =* 0.487). Dashed gray lines give global means across all families for each trait.

**Figure 5.**
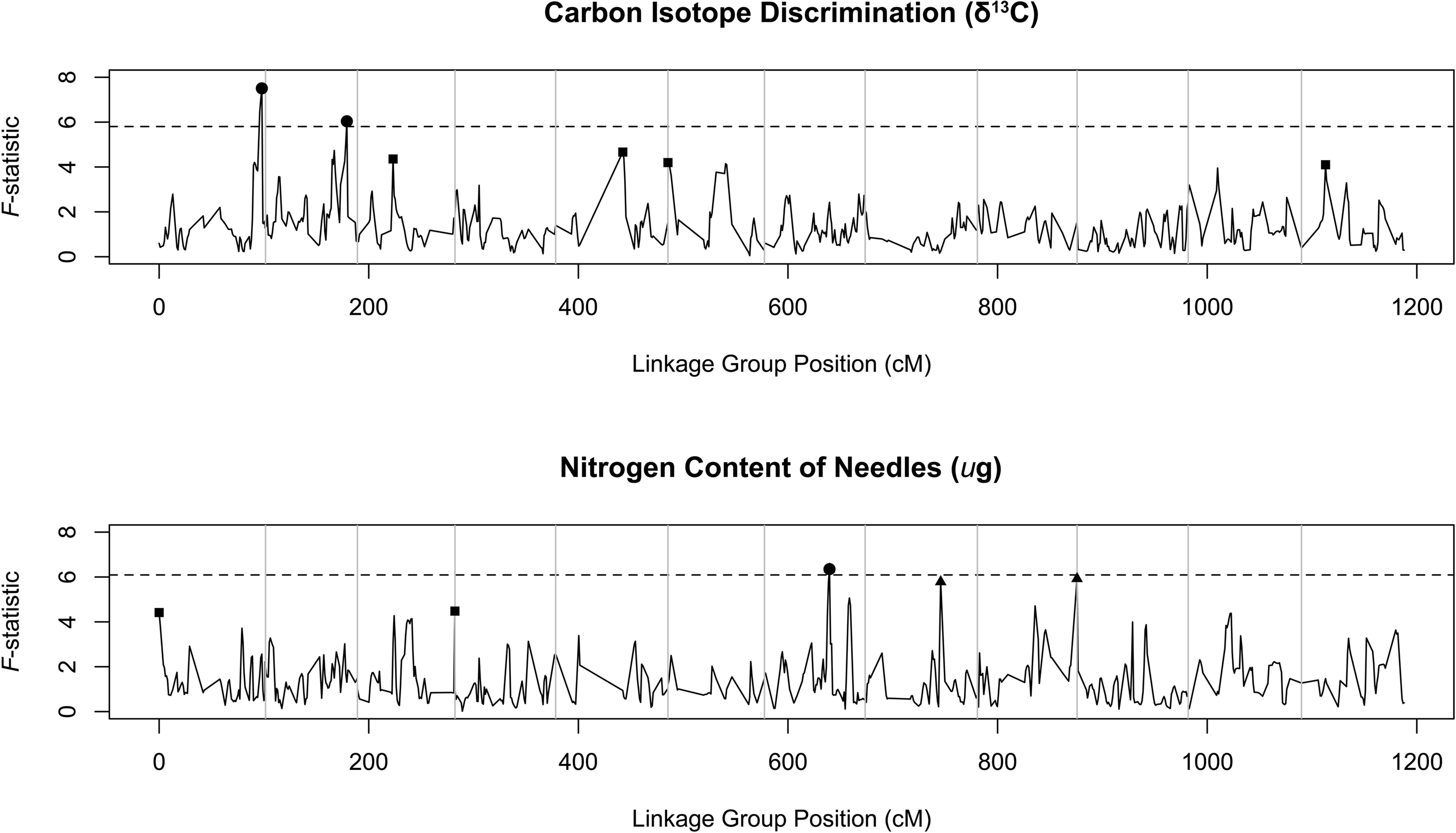
The distributions of the *F*-statistic derived from single QTL models across each linkage group for carbon isotope discrimination and nitrogen content of needles reveals the isolated nature of QTLs. The dashed horizontal line in each panel is the genome-wide significance threshold (α = 0.05) for the *F*-statistic based on the permutation scheme (*n* = 1,000 permutations) suggested by Churchill and Doerge (1994). Significant QTLs are denoted with filled circles (α = 0.05, genome-wide), filled triangles (α = 0.01, chromosome-wide) or filled squares (α = 0.05, chromosome-wide).

**Table 5.**
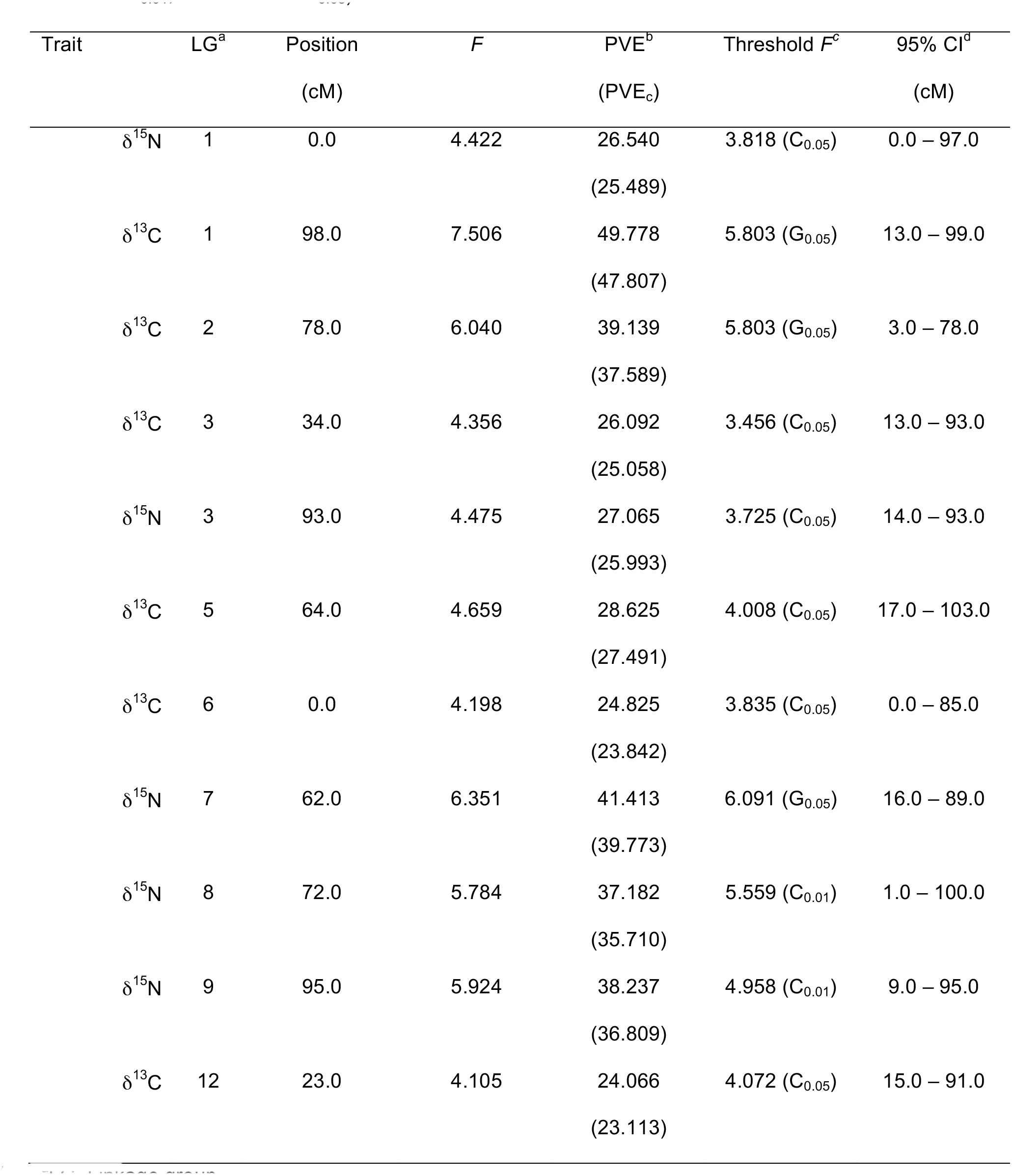
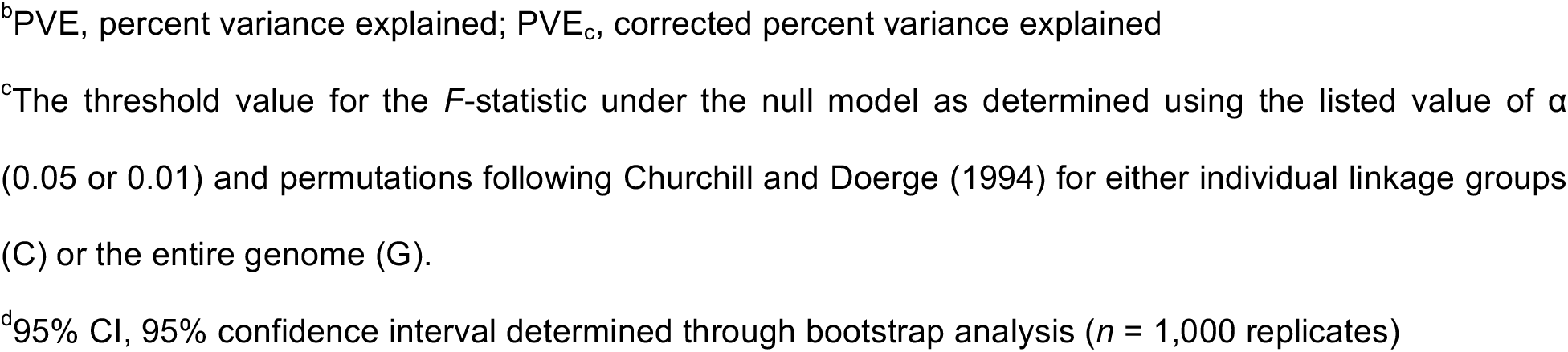
Summary of QTLs for each trait that survive multiple test corrected significance thresholds at either the level of the whole genome (α = 0.05 for G_0.05_) or a chromosome (α = 0.01 for C_0.01_, α = 0.05 for C_0.05_)

| Trait | LG<sup>a</sup> | Position<br>(cM) | *F* | PVE<sup>b</sup><br>(PVE<sub>c</sub>) | Threshold *F*<sup>c</sup> | 95% CI<sup>d</sup><br>(cM) |
| --- | --- | --- | --- | --- | --- | --- |
| $\delta^{15}\text{N}$ | 1 | 0.0 | 4.422 | 26.540<br>(25.489) | 3.818 ($C_{0.05}$) | 0.0 – 97.0 |
| $\delta^{13}\text{C}$ | 1 | 98.0 | 7.506 | 49.778<br>(47.807) | 5.803 ($G_{0.05}$) | 13.0 – 99.0 |
| $\delta^{13}\text{C}$ | 2 | 78.0 | 6.040 | 39.139<br>(37.589) | 5.803 ($G_{0.05}$) | 3.0 – 78.0 |
| $\delta^{13}\text{C}$ | 3 | 34.0 | 4.356 | 26.092<br>(25.058) | 3.456 ($C_{0.05}$) | 13.0 – 93.0 |
| $\delta^{15}\text{N}$ | 3 | 93.0 | 4.475 | 27.065<br>(25.993) | 3.725 ($C_{0.05}$) | 14.0 – 93.0 |
| $\delta^{13}\text{C}$ | 5 | 64.0 | 4.659 | 28.625<br>(27.491) | 4.008 ($C_{0.05}$) | 17.0 – 103.0 |
| $\delta^{13}\text{C}$ | 6 | 0.0 | 4.198 | 24.825<br>(23.842) | 3.835 ($C_{0.05}$) | 0.0 – 85.0 |
| $\delta^{15}\text{N}$ | 7 | 62.0 | 6.351 | 41.413<br>(39.773) | 6.091 ($G_{0.05}$) | 16.0 – 89.0 |
| $\delta^{15}\text{N}$ | 8 | 72.0 | 5.784 | 37.182<br>(35.710) | 5.559 ($C_{0.01}$) | 1.0 – 100.0 |
| $\delta^{15}\text{N}$ | 9 | 95.0 | 5.924 | 38.237<br>(36.809) | 4.958 ($C_{0.01}$) | 9.0 – 95.0 |
| $\delta^{13}\text{C}$ | 12 | 23.0 | 4.105 | 24.066<br>(23.113) | 4.072 ($C_{0.05}$) | 15.0 – 91.0 |

<sup>a</sup>LG, Linkage group
<sup>b</sup>PVE, percent variance explained; PVE<sub>c</sub>, corrected percent variance explained
<sup>c</sup>The threshold value for the *F*-statistic under the null model as determined using the listed value of $\alpha$ (0.05 or 0.01) and permutations following Churchill and Doerge (1994) for either individual linkage groups (C) or the entire genome (G).
<sup>d</sup>95% CI, 95% confidence interval determined through bootstrap analysis ( $n = 1,000$ replicates)

For the 11 QTLs detected using one-locus models, 10 were consistent with multiple QTLs using two-locus models (Table 6). In general, the QTLs from the one-locus models were one of the pair of QTLs detected in the two-locus models. There were four exceptions to this pattern, with two of these exceptions being a minor modification in position of the original QTL equal to 1.0 cM. The other two exceptions included significant changes to the position of the original QTL, with the QTL on linkage group 3 for δ^15^N changing from 93.0 cM to 52.0 cM and 35.0 cM and the QTL on linkage group 6 for δ^13^C changing from 0.0 cM to 46.0 cM and 56.0 cM (Tables 5 and 6). The average spacing between QTLs on the same linkage group was 29.4 cM, with a minimum of 3 cM to a maximum of 85 cM. The multi-QTL PVE for each trait ranged from a minimum of 42.685% to a maximum of 71.315%, with only one instance of positional overlap in QTLs for each trait (linkage group 3 at 34.0 cM for δ^13^C and 35.0 cM for δ^15^N). On average, there was a negative relationship between distance (cM) and the correlation of family effects (Pearson’s *r*) between QTLs on the same linkage group (Figure S18), so that strong positive correlations of family effects were observed when QTLs were close together (<15 cM) and strong negative correlations when QTLs were farther apart (>20 cM).

**Table 6.** Summary of two QTL models fit to each significant QTL from Table 4. Bolded *P*-values are less than 0.05.

| Trait | LG<sup>a</sup> | Position 1<br>(cM) | Position 2<br>(cM) | *F* | *P* | PVE<sup>b</sup> |
| --- | --- | --- | --- | --- | --- | --- |
| $\delta^{15}\text{N}$ | 1 | 0.0 | 79.0 | 3.89 | **0.0023** | 54.518 |
| $\delta^{13}\text{C}$ | 1 | 98.0 | 13.0 | 2.92 | **0.0149** | 64.725 |
| $\delta^{13}\text{C}$ | 2 | 77.0 | 66.0 | 3.76 | **0.0030** | 61.685 |
| $\delta^{13}\text{C}$ | 3 | 34.0 | 14.0 | 3.18 | **0.0091** | 44.459 |
| $\delta^{15}\text{N}$ | 3 | 52.0 | 35.0 | 4.24 | **0.0012** | 57.594 |
| $\delta^{13}\text{C}$ | 5 | 64.0 | 88.0 | 1.81 | 0.1135 | 37.745 |
| $\delta^{13}\text{C}$ | 6 | 46.0 | 56.0 | 3.84 | **0.0026** | 48.892 |
| $\delta^{15}\text{N}$ | 7 | 62.0 | 80.0 | 4.69 | **0.0005** | 71.315 |
| $\delta^{15}\text{N}$ | 8 | 71.0 | 68.0 | 2.57 | **0.0287** | 49.661 |
| $\delta^{15}\text{N}$ | 9 | 95.0 | 64.0 | 2.90 | **0.0155** | 53.602 |
| $\delta^{13}\text{C}$ | 12 | 23.0 | 43.0 | 3.20 | **0.0088** | 42.685 |

<sup>a</sup>LG, Linkage group
<sup>b</sup>PVE, percent variance explained by both QTLs

QTL effects from the one-locus QTL models were consistent with differentiation between regional populations, with family effects opposite in sign more often than expected by chance for δ^13^C (Fisher’s exact test: odds ratio = 0.113, *P* = 0.009), but not for δ^15^N (Fisher’s exact test: odds ratio = 1.319, *P* = 1.0). Trait differentiation was similarly structured (Tables 3 and 4), with the clearest signal of differentiation for δ^13^C. The same patterns were observed for family effects in the two-locus models for the original QTL from Table 5, but not for the second QTL (*P* > 0.05 for both δ^13^C and δ^15^N).

## Discussion

Climate is one of the main drivers for the distribution and diversification of forest tree species (MacArthur 1972; Royce and Barbour 2000; Ettinger *et al*. 2011; Alberto *et al*. 2013). The relative importance of specific climate variables as drivers of natural selection, however, is often assumed. For example, if a phenotypic trait is correlated to water availability in one species, the same trait is often studied in a different focal species without documenting water availability as having a large impact on fitness variation in the latter. The problem lies in the assumption that this correlation is also indicative of similar fitness consequences across species. Here, we address this issue for foxtail pine using a novel combination of species distribution modeling and quantitative genetics. We illustrate the importance of water availability to the distribution of foxtail pine and hence fitness, as well as describe the genetic architecture of WUE, a phenotypic trait responsive to water availability, so that this trait and the markers correlated to it can be used to test hypotheses about local adaptation and its genetic architecture.

### Climate drivers of the current geographical distribution and WUE

In many situations, drivers of geographical distributions for tree species are obvious. For example, links between light availability, temperature, precipitation, and phenological traits are commonly noted for forest trees (Howe *et al*. 2003; Chuine 2010). In other situations, however, climate drivers are less clear, so that quantification of the relative importance for a suite of climate variables is needed. For foxtail pine, the drivers of its current geographical distribution appear to be a mixture of temperature-related and precipitation-related variables, with a clear pattern that precipitation-related variables are necessary to explain the current geographical range. This implies that phenotypic traits correlated to precipitation-related variables likely have fitness consequences for foxtail pine, as precipitation-related variables appear to structure its current range. Additionally, the importance of these drivers is differentiated between regional populations, with precipitation-related variables more differentiated than temperature-related variables, which mimicked differentiation of phenotypic trait values. Thus, if we leverage the correlations between δ^13^C and water availability, a crucial component of survival and hence fitness, observed in other plant species (Ehleringer *et al*. 1993) and the conclusion that precipitation-related variables are important for the distribution of foxtail pine, it is likely that δ^13^C variation in foxtail pine is linked with fitness.

In general, increases in δ^13^C reflect higher WUE (Farquhar *et al*. 1982). Inspection of mean values for δ^13^C for each region (see Figure 3), in light of the documented precipitation patterns, however, appears contradictory. On average, maternal trees in the Klamath Mountains had higher δ^13^C values, which suggests higher WUE, yet precipitation is much higher in the Klamath Mountains than in the southern Sierra Nevada. It is well known, however, that soil properties, such as coarseness and depth to bedrock, affect available soil moisture. For example, small differences in soil texture observed across the Southern Sierra Nevada Critical Zone Observatory, a site not far removed from the regional population of foxtail pine in the southern Sierra Nevada, result in large differences in the available soil moisture (Bales *et al*. 2011). Soil texture also varied by elevation, with soils at the highest elevations being coarser and less developed. As such, water availability in these soils was more limited even though snowfall was typically higher. Soils between regional populations of foxtail pine are fundamentally different, and so is the local distribution of foxtail pine. In the Klamath Mountains, soils are primarily ultramafic, while in the southern Sierra Nevada they are largely granitic. Foxtail pine grows near tops of local peaks in the Klamath Mountains, whereas in the southern Sierra Nevada it is distributed broadly across large swathes of high elevation sites. Thus, one explanation for the apparent contradiction is that soil properties are different, so as to create patterns of soil moisture not reflective of regional mean precipitation patterns. Foxtail pine in the Klamath Mountains often inhabits areas with high levels of boulder cover (Eckert and Sawyer 2002; Eckert 2006), which are expected to house soils with less capacity to hold water over long periods of time. When coupled with the higher average temperatures in the Klamath Mountains, this suggests that water may be more limited throughout the year (e.g. summer drought) than expected based on annual precipitation totals. Additional work, however, would be needed to quantify trait variation within each regional population and correlate it to both climate and soil characteristics.

### Genetic architecture of water-use efficiency

Both δ^13^C and δ^15^N were consistent with non-zero heritabilities. Families and regions accounted for approximately 50% of the total phenotypic variance for δ^13^C and 30% for δ^15^N. Models with effects due to families or families nested within regions were also strongly preferred over models without these effects (Table 4). The effect of region, however, was highest in magnitude for δ^13^C, with the variance component for region larger than that for family. This is consistent with previous estimates of quantitative genetic parameters for these phenotypic traits in other conifers. For example, δ^13^C and δ^15^N are both heritable in a variety of pine species (Brendel *et al*. 2002; Baltunis *et al*. 2008; Gonzalez-Martinez *et al*. 2008; Cumbie *et al*. 2011; Joao Gaspar *et al*. 2013; Marguerit *et al*. 2014; Eckert *et al*. 2015). Populations within many species are also often differentiated for δ^13^C, but not for δ^15^N (e.g. Eckert *et al*. 2015; Maloney *et al*. unpublished). Further work, however, would be needed to precisely estimate the level of differentiation for these traits, as well as to test whether this level of differentiation is larger than that expected for neutral loci (i.e. this pattern is consistent with local adaptation).

Estimates of narrow-sense heritabilities (*h*^2^) resulted in values greater than 1.0 for each phenotypic trait no matter which model with a family effect was used (i.e. families or regions plus families nested within regions). This could be due to tissue sampling occurring prior to formation of randomized blocks in the common garden, as family groups would be confounded with micro-environmental variation. Use of data from Eckert *et al*. (2015) and Maloney *et al*. (unpublished data) for sugar pine (*P. lambertiana* Dougl.), western white pine (*P. monticola* Dougl.), and whitebark pine (*P. albicaulis* Engelm.) grown at the same facility in the same experimental conditions, however, reveals that block effects for δ^13^C were present only for the relatively fast growing western white pine (Type III Wald *F*-tests with Kenward-Rogers degrees of freedom; sugar pine: *F*_1,416.49_ = 3.5166, *P* = 0.06146; western white pine: *F*_1,630.24_, *P* = 0.00068; whitebark pine: *F*_1,452.75_ = 0.0147; *P* = 0.9037). In contrast, block had a statistically significant effect on δ^15^N for sugar pine and western white pine (*P* < 0.001), but not whitebark pine (*F*_1,429.22_ = 1.6252, *P* = 0.20305). Thus, our results should be taken with caution, but family effects estimated here were similar in magnitude to those from Eckert *et al*. (2015) and randomized blocks tended to have no effect on the same phenotypic traits measured in whitebark pine at the same facility, a species with a similar pattern of early slow growth (McCune 1988).

If our results are indicative of true signal, effect sizes could be over-estimated on average due to the small number of sampled families (Beavis 1994). To illustrate this effect, we re-analyzed the data from Eckert *et al*. (2015) for sugar pine, which was grown in a common garden at the same facility and measured for δ^13^C using the same methodology, by resampling smaller numbers of families (*n* = 108 families resampled in decreasing numbers from 108 to three families) and estimating *h*^2^. As the number of sampled families decreased, estimates of mean *h*^2^ became larger (Figure S19), with a 1.5-fold increase in the mean *h*^2^ as the number of sampled families dropped from 108 to three. This is likely also the case for foxtail pine and for δ^15^N. Regardless of the precise value of *h*^2^, it is clear that at least a moderate amount of segregating genetic variation exists for this trait in natural populations of foxtail pine.

There was also a moderate, but statistically insignificant, positive correlation between δ^13^C and δ^15^N (Figure 4). This has been noted in other species, such as loblolly pine (Cumbie *et al*. 2011), although general patterns in the sign of the correlation are lacking. In this context, positive correlations could indicate that WUE is determined primarily through leaf-level assimilation (e.g. Johnson *et al*. 1999; Prasolova *et al*. 2005), while a negative correlation could indicate that WUE is determined primarily through stomatal conductance. Despite the observed positive correlation, little evidence of pleiotropy was detected, with only a single QTL on linkage group 3 shared between traits. The lack of pleiotropy for these traits has been noted in several other conifer species (e.g. Marguerit *et al*. 2014). Correlations between δ^13^C and δ^15^N, or growth traits more generally, can also be driven environmentally and can change depending on water availability. For example, Joao Gaspar *et al*. (2013) have shown that in water limiting environments δ^13^C correlates with survival, but in less water limited environments δ^13^C correlates with height growth for maritime pine (*P. pinaster* Ait.). A similar case might be occurring for foxtail pine, where in the wetter Klamath Mountains δ^13^C variation is correlated with overall growth and in the more xeric southern Sierra Nevada it is correlated with survival. In this context, WUE would be realized through leaf-level assimilation in the Klamath region (as in Weih *et al*. 2011 for *Salix*), and through stomatal conductance in the southern Sierra Nevada. Sampling more families, measurement of other traits (e.g. growth), and experimentation in multiple environments, however, would be needed to test these ideas. Importantly, δ^13^C should be measured within natural populations to assess correspondence between inferences from common gardens and natural populations.

Using one-locus QTL models, the observed segregating genetic variance for δ^13^C was dissected into two major QTLs and four suggestive QTLs (Table 5). Each QTL explained a large fraction of total phenotypic variance (23.113% to 47.807%), which suggests that the genetic architecture of this fitness-related trait includes loci of large effect. Under many models of adaptation, however, is difficult to separate QTLs composed of a single, large-effect locus from those composed of several small-effect loci (Yeaman and Whitlock 2011). The observed large values of PVE may also be over-estimated (Beavis 1994), although there is precedence for large effect QTLs for δ^13^C in other species of *Pinus*, especially those distributed in water-limited regions displaying moderate levels of genetic differentiation among populations. For example, Marguerit *et al*. (2014) identified a QTL explaining 67% of phenotypic variance for δ^13^C in maritime pine, which is distributed across the Mediterranean regions of Europe and has moderate levels of genetic structure across this range (Eveno *et al*. 2008). For foxtail pine, water availability is an important driver of its current geographical distribution and genetic structure is moderate to high between regional populations and among stands within regional populations (Eckert *et al*. 2008, but see Oline *et al*. 2000). Furthermore, family effects for these QTLs were consistent with differentiation among regions, so it is plausible that the architecture discovered here for δ^13^C largely represents genomic regions underlying trait divergence between the regional populations. If this is the case, this architecture has evolved since the divergence of the regional populations from their common ancestor on the order of one million years ago (Eckert *et al*. 2008).

Summaries of the results from two-locus QTL models were largely consistent with those from the one-locus models. For the 11 QTLs reported in Table 5, 10 were consistent with at least two segregating QTLs. This brings the total number of QTLs to four major and seven suggestive QTLs for δ^13^C and two major, four minor, and four suggestive QTLs for δ^15^N. Interestingly, the correlation of family-level effects for the two QTLs on the same linkage group was negatively related to the distance between these QTLs, so that QTLs close together tended to have similar patterns of family-level effects, whereas QTLs at larger distances tended to have opposite family-level effects (Figure S14). This trend was uncorrelated with the difference in effect sizes between QTLs. When added to the observation that family effects were often consistent within regions and differentiated between regions, a likely explanation for this pattern is some form of natural selection driving clustering of loci dependent on consistency of their effects on a fitness-related trait. The fitness benefit of clustering, however, is related to the level of gene flow (Yeaman and Whitlock 2011), so that clustering of adaptive alleles is expected under high levels of gene flow, reduced recombination, and strong magnitudes of selection. This is especially pronounced when genomic rearrangements are common. Inspection of the family-level linkage maps from Friedline *et al*. (2015), however, reveled little evidence for clustered QTLs displaying differing marker orders across families more so than random positions on the linkage map. This explanation, however, is complicated given that gene flow is approximately zero between these regions (Eckert *et al*. 2008) and populations of foxtail pine are unlikely to be at selection – migration equilibrium due to large effective population sizes and long generation times. For example, patterns of segregating ancestral variation after divergence are similar to those predicted by gene flow (Pamilo and Nei 1988), so that it becomes difficult to separate pattern from process with regard to the effects of gene flow on adaptive genetic architectures. Additional work within natural populations, including fine mapping of trait values in the linkage bins defined by Friedline *et al*. (2015), would be needed to test these ideas further.

We leveraged the annotations of contigs at or near (± 3 cM) the estimated QTL positions to search for putatively functional genes as the drivers of the genotype-phenotype correlations for each QTL (Table S3). Annotations for foxtail pine contigs were derived through similarity searches against the loblolly pine genome. Annotations were obtained from any locus on a loblolly pine scaffold containing a significant hit to a RADtag from foxtail pine, with significance justified by the estimated substitution rate and divergence time between these species (Friedline *et al*. 2015). Several statistically significant QTLs had no annotation information available. For example, the QTL on linkage group 1 for δ^13^C had no annotations available within a 6-cM window encapsulating the QTL, despite 24 of 76 RADtags having significant similarity to scaffolds in loblolly pine. This is consistent with reports of gene densities reported for conifers (Nystedt *et al*. 2013; Neale *et al*. 2014). For the QTL related to δ^13^C on linkage group 2 (Table 5), however, two of the 18 RADtags for foxtail pine had sequence similarity to loblolly pine scaffolds, with annotated InterPro domains suggestive of loci encoding stress responsive proteins (Table S3; Toka *et al*. 2010; Karijolich *et al*. 2015). Another example of potentially biologically informative results included the QTL on linkage group 9 for δ^15^N where putative homologs for proteins with domains such as ribosomal protein L38e, cytochrome P450, and thiolase were present. Proteins containing these domains have been implicated in lipid turnover during leaf senescence (Troncoso-Ponce *et al*. 2013), as well as plant growth and drought stress response (Tamiru *et al*. 2015). Care should be taken in interpreting these results, however, as QTL intervals were wide, annotations were based on statements of homology with gene predictions in an early release of the loblolly pine genome sequence (Wegrzyn *et al*. 2014), and *post hoc* explanations linking gene products to phenotypic traits is prone to storytelling (Barrett and Hoekstra 2011; Pavlidis *et al*. 2012). It is important to note, however, that these concerns are with interpretations of putative functions of genes located within the QTL as sensible in their effect on the measured phenotypic trait, and not with the biological signal of linkage driving the discovery of the QTL.

## Conclusions

We have used a mixture of species distribution modeling and quantitative genetics to test two hypotheses about WUE, as measured by δ^13^C, for foxtail pine. We showed that precipitation-related variables structured the geographical range of foxtail pine, that climate-based niches differed between regional populations, and that similar patterns were apparent for δ^13^C, which was also demonstrated to be heritable. We subsequently dissected this heritability into a set of large-effect QTLs (*n* = 21 total, with 11 for δ^13^C and 10 for δ^15^N), which we interpret in light of population genetic theory about local adaptation. While we cannot definitely say that WUE, as measured by δ^13^C, contributes to local adaptation, we have described to a first approximation its genetic architecture, while noting several patterns consistent with δ^13^C being a fitness-related trait affected by natural selection. These are useful results with which to generate further hypotheses about the evolution of genetic architecture contributing to local adaptation in natural populations (e.g. Holliday *et al*. 2015). Our results also shed light on ecologically relevant phenotypic trait variation useful for management decisions and predictions for range shifts under changing climates.

## Acknowledgements

We are grateful to the staff at the USDA Institute of Forest Genetics for space and logistical support in the establishment of the common garden, as well as the staff at the VCU Nucleic Acids Research Facility and Center for High Performance Computing for help in generating and analyzing the Illumina short read data. Seeds for foxtail pine were obtained from Tom Blush and Tom Burt. Rodney Dyer, Christopher Gough, William Eggleston and Salvatore Agosta all made helpful comments on D. E. Harwood’s M.S. thesis at VCU, which was the basis for this work.

## Data Archiving Statement

- **Genotype data** are available as raw short read data as part of NCBI BioProject PRJNA310118, processed short read data in VCF format (File S1), and imputed data in a tab-delimited text file (File S2).
- **Phenotypic trait data** are available for all half-siblings within each of the five families used for QTL mapping in a tab-delimited text file (File S3).
- **Location data** used for species distribution modeling are available in a tab-delimited text file (File S4).

## Compliance with Ethical Standards

- **Disclosure of potential conflicts of interest:** Funding for this work was provided from Virginia Commonwealth University (start-up funds awarded to A. J. Eckert) and from the National Science Foundation (NSF-NPGI-PRFB-1306622 awarded to C. J. Friedline).
- **Research involving Human Participants and/or Animals:** There were no human participants or animals used in this research.
- **Informed consent:** There were no human participants used in this research, so informed consent is not applicable.

